# A Holistic Analysis of the Intestinal Stem Cell Niche Network

**DOI:** 10.1101/2019.12.12.871756

**Authors:** Darrick M. Hansen, Paloma Ivon Meneses Giles, Xi C. He, Shiyuan Chen, Ariel Paulson, Christopher M. Dekaney, Jennifer Wang, Deqing Hu, Aparna Venkatraman, Woosook Kim, John Kaddis, Barbara J. Olack, James C.Y. Dunn, Calvin Kuo, Susan Henning, Alan M. Hanash, Courtney W. Houchen, John Lynch, Martin G. Martin, Joyce C. Niland, Matthias Stelzner, Melissa Wong, Timothy C. Wang, Jian Yu, Kelley Yan, Linheng Li

## Abstract

Although many studies into the intestinal stem cell (ISC) niche have been carried out, they have focused on the role of a single cell type or molecular signal. However, no holistic comparisons of the predominant cell types and signals present within the intestinal mucosa have been conducted to date. We utilize bulk RNA sequencing to profile 20 different mucosal cell types covering four major cell categories: epithelial, stromal, endothelial and immune. We further examined the stromal signaling environment using scRNAseq to provide a more comprehensive view of the signaling microenvironment within the intestinal mucosa. We identified the primary signals for the major ISC regulatory pathways and their respective cellular sources. Our analysis suggests that a ‘niche network’ exists, with no single cell type being responsible for ISC self-renewal, proliferation, or differentiation; rather, each cell type within the network carries out specific functions in a highly cooperative and coordinated manner.

## Introduction

The intestine is arguably the most multifunctional organ within the body, representing a remarkable intersection of some of the most complex biological systems known: the epithelium serves as a highly selective barrier, the vasculature nourishes the tissue while transporting vital nutrients away, the immune system must fend off pathogens without attacking food material, while the stroma serves as a scaffold and signaling relay. These cooperative functions are likely dependent on an exquisitely coordinated signaling network, in which intestinal stem cells (ISCs) take a central position.

ISCs reside in specialized compartments known as crypts, where they generate multipotent progenitors called transit-amplifying cells (TACs) which undergo rapid division. TACs differentiate into a variety of cell lineages including enterocytes, enteroendocrine cells, Goblet cells, and Paneth cells. While Paneth cells remain at the crypt bottom and interdigitate with ISCs, the remaining cell types migrate upward along the crypt-villus axis as they differentiate into their various specialized roles (Sailaja et al., 2016). Under homeostatic conditions, actively cycling ISCs are responsible for the repopulation of the entire intestinal epithelium; In the event of injury however, slow-cycling reserve ISCs (rISCs) mediate tissue repair with assistance from a subset of differentiated cells which reacquire their ‘stem’ state through mechanisms of cellular plasticity (Li and Clevers, 2010, Scoville et al., 2008, Yousefi et al., 2017). The collection of structural, molecular, and cellular components within the stem cell’s immediate vicinity are collectively known as the niche, and changes in any of these components will alter a stem cell’s behavior to better meet the body’s needs. ISC selfrenewal, cell cycle state, metabolic activity, lineage specification, and many other important processes in stem cells are all controlled through niche dynamics (Sailaja et al., 2016).

The intestinal epithelium experiences some of the harshest and most variable conditions within the body, all while maintaining its highest proliferative rate (Barker, 2014). Such variability likely necessitates a very responsive and complicated network of molecular signals generated by many different cells. Though previous attempts to dissect the ISC niche have made remarkable progress, they have generally employed a one-against-all analytical approach (He et al., 2004, San Roman et al., 2014, Stzepourginski et al., 2017, Worthley et al., 2015, Aoki et al., 2016, Sato et al., 2009) where a single cell type or signal is compared to the ‘other’ homogenized components. Such approaches likely oversimplify an intricate network and dilute out rare or highly specific components within the ‘other’ category. Furthermore, any direct interactions with another unexamined cell type will likely be missed, making it difficult to draw conclusions about whether the observed effects on ISCs are carried out through direct interactions or indirectly through one or multiple intermediate cell types. A catalog of the predominant molecular, cellular, and structural components of the intestinal mucosa would increase researchers’ awareness of considerations outside their area of expertise and help them prioritize their search for potentially confounding variables. Therefore, a systematic investigation of the major cellular and molecular components of these structures and how their organization might govern tissue function is key to understanding the ways aberrant conditions affect or are affected by this complex milieu.

To gain a more complete understanding of the predominant factors within the intestinal microenvironment under homeostatic conditions, 20 cell types belonging to one of four categories: epithelial, stromal, endothelial, and immune, were collected. RNA-Sequencing analysis was then conducted on both bulk (bRNA-Seq) and single cell (scRNA-Seq) samples, and our results were confirmed using fluorescence imaging. The major signals identified were then catalogued, their cellular sources compared, and their spatial distribution determined.

## Results

### Characterization of Cell Types and Categories

In order to gain a more comprehensive understanding of the relative contribution these cell categories and types make to the major signaling networks within the intestinal mucosa, we first used bRNA-Seq to profile key cell types within the epithelial, endothelial, stromal, and immune categories. To be able to reflect more faithfully the spatial distribution of cells within the intestinal mucosa, we have adapted previously established 3D immunofluorescent staining and imaging techniques (Bernier-Latmani and Petrova, 2016, Appleton et al., 2009). The longitudinal view clearly lays out the smooth muscle as forming the base for the crypt and villus structures (Fig. 1A-E). Epithelial cells form a monolayer which adheres to the lamina propria via a basement membrane which is contiguous with the basement membrane separating the mucosa from the muscularis (Fig. 1A-B). Endothelial cells form a mesh-like network that is distributed throughout the stromal compartment which establishes a tight association with the basement membrane of the villus (Fig. 1C). CD45+ immune cells are sporadically distributed throughout the mucosa (Fig. 1D). A complete list of all 20 cell types from the 4 distinct categories along with their isolation methods and descriptions can be found in Supplementary Figure 1A.

**Figure 1.**
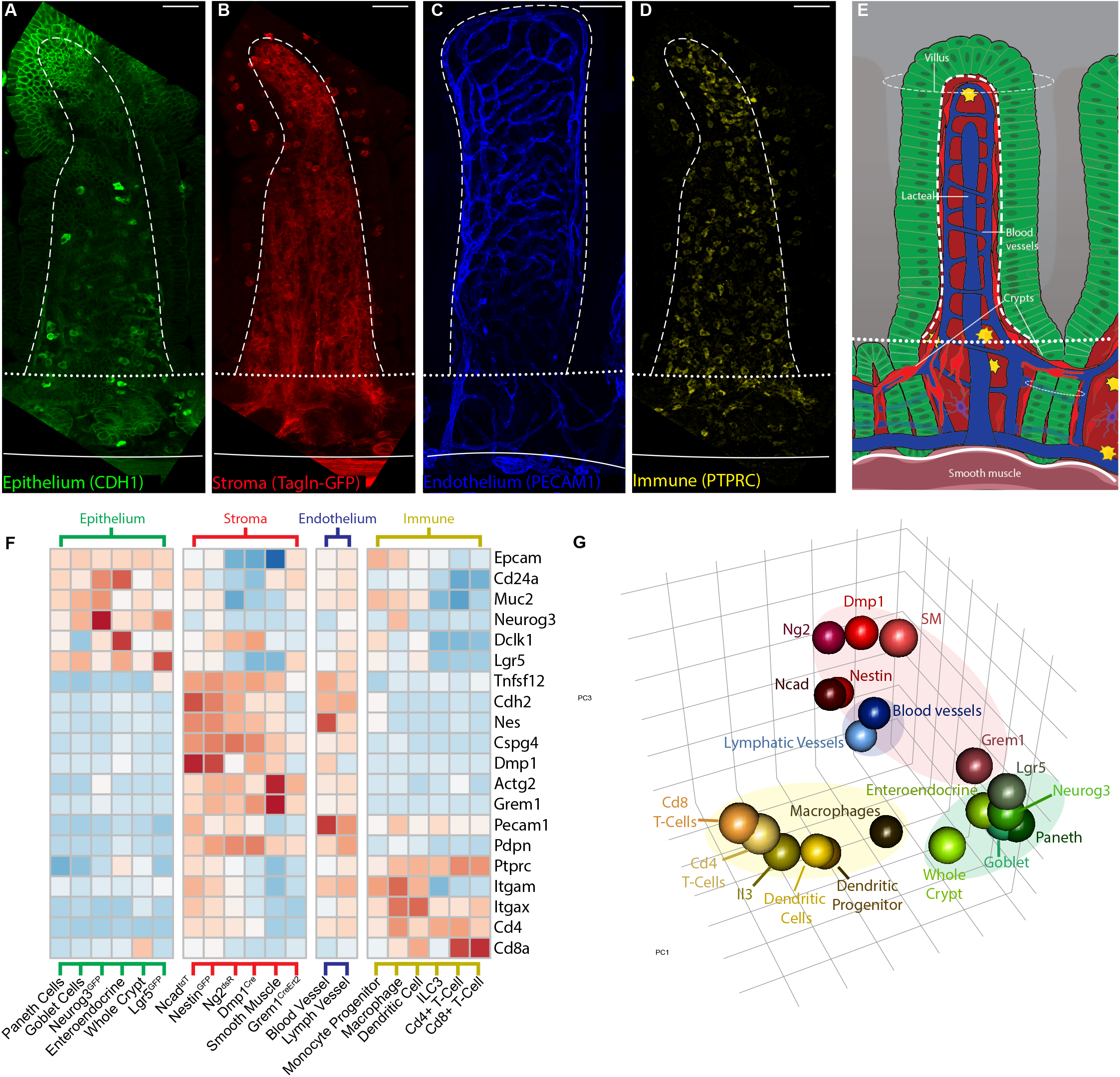
Characterization of the Cell types and compartments examined. (A-E) representative images showing the juxtaposition and spatial relationships between the various cell categories and regions examined. (F) Centered and scaled expression analysis of the genes used for cell type and category identification. (G) Principal Component analysis of the 20 cell types used in bulk sequencing Solid lines mark the muscularis/mucosa border, dotted lines mark the border of the villus/pericryptal regions, and dashed lines mark the stromal/epithelial border. Scale bars, 50μm See Figure S1 for a complete list and details of the cell types used along with the functional genes used to confirm their identities.

While immunostaining was used to identify epithelial, endothelial, and immune cell types, stromal populations were assigned to the genetic markers used to isolate them. The stroma of the intestinal mucosa has received substantially less attention than the other cell categories; few functionally distinct cell types have been defined and fewer still have had their roles within the ISC niche examined. We therefore selected genetic models that have been used to characterize the stem cells and niches of other tissues in order to determine whether they perform similar functions within the small intestine. The five mouse models we used to profile the stroma were: Grem1^CreErt2^+, which has been shown to mark multipotent skeletal stem cells (Worthley et al., 2015); Ncad^tdT^+ mice have been used to characterize the hematopoietic stem cell (HSC) niche and how *Cdh2*+ cells steer HSCs to a quiescent or active state (Zhao et al., 2019); both Nestin^GFP^+ and Ng2^dsR^+ models have been used to study neural stem cells and how mural cells of the bone marrow influence HSC function (Mignone et al., 2004, Zhu et al., 2008, Sa da Bandeira et al., 2017, Mendez-Ferrer et al., 2010); lastly is the Dmp1^Cre^+ model, which has been shown to mark cells that guide both mesenchymal stem cell (MSC) and HSC activity, in addition to overlapping extensively with PDGFRA^high^ subepithelial cells which are known to be important regulators of ISC function (Lim et al., 2017, Shoshkes-Carmel et al., 2018). One notable categorical omission from our data is the enteric nervous system. Neurons have been demonstrated to play important roles in ISC regulation and intestinal regeneration, however we were unable to obtain data of sufficient quality from intestinal neurons to include in this study.

Expression of the genes used in FACS isolation showed the expected compartmental and cell type specificity (Fig. 1F). *Cdh2* (encoding N-cadherin) showed highest expression levels in Ncad^tdT^+ cells while reaching intermediate levels in Nestin^GFP^+ cells, with the reciprocal being true for *Nes* expression. These data confirm the quality of the collection and analysis techniques employed overall. The expression values of the genes used in FACS isolation were not always highest in their respective cell type however, which is likely due to the Cre system being a qualitative measure at a distinct temporal point rather than quantitative. *Dmp1* is one example, with expression that was higher in Nestin^GFP^+ and Ncad^tdT^+ cells than in Dmp1^Cre^+ derived cells.

As expected, principle component analysis of the various transcriptomes resulted in the cell types clustering according to their categorical distinctions (Fig. 1G). To briefly summarize, Grem1^CreErt2^+ cells adopted a unique position between stromal and epithelial cells. Nestin^GFP^+ and Ncad^tdT^+ cells clustered together, consistent with their overlap and distribution pattern. Intriguingly, Ng2^dsR^+ and Dmp1^Cre^+ cells were transcriptomically similar, yet showed differences in several genes important to cellular function (Fig. S1B). Innate lymphoid type 3 cells (ILC3) were positioned between adaptive T Cells and Innate Immune Dendritic cells and Macrophages (Fig. 1G).

### Characterization of Genetic Models and Stromal Populations

Both Nestin^GFP^+ and Ncad^CreErt2^+ stromal cells were present from the smooth muscle to the villus tips (Fig. 2A-B) and possessed somewhat similar distributions and morphologies to the niche cells described within the bone marrow. Though there was considerable overlap and similarities between these genetic markers, several differences were apparent. Ncad^CreErt2^+ cells were more likely to encircle the vasculature with a flattened morphology, while Nestin^GFP^+ cells frequently laid on top of these cells and possessed long filopodia which connect with one another (Fig. S2A-E) Ncad^CreErt2^+ cells were more likely to be present at the lower “ring”, while Nestin^GFP^+ were more likely to localize to the isthmus. Ng2^dsR^+ cells also overlapped with Nestin^GFP^+ cells but were the only cell type examined that exhibited a strong spatial bias within the mucosa, being almost exclusively found within the villi (Fig. 2C,S2B). Unfortunately, their morphological characteristics could not be determined due to the tendency of the dsRed protein to form aggregates, preventing even cytosolic distribution.

**Figure 2.**
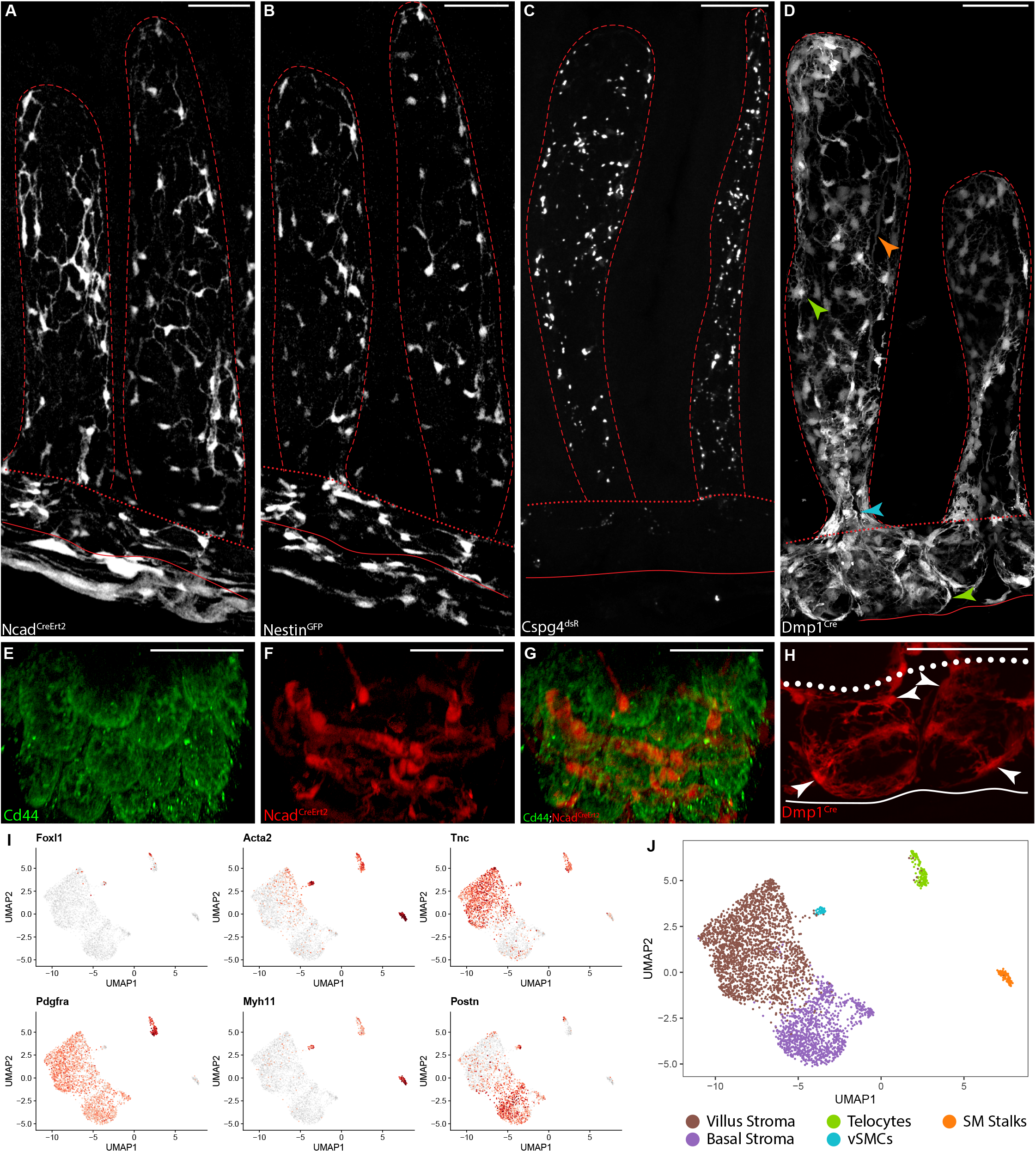
Characterization of the Genetic Models and Stromal Cell Types within the lamina propria. (A-D) Representative images of the distribution of genetic markers within the small intestine. (E-G) Whole mount images of Ncad^CreErt2^+ cells and crypt bases (CD44) demonstrating the lower “ring” structures found encircling the crypts. (H) Whole mount images of Dmp1^Cre^+ telocytes illustrating the typical “+4” and isthmus localization (I-J) scRNA-Seq analysis of the marker genes used to determine cluster identities and the assigned names. Solid lines mark the muscularis/mucosa border, dotted lines mark the border of the villus/pericryptal regions, and dashed lines mark the stromal/epithelial border. Scale bars, 50μm See Figure S2 for a more detailed view of the pericryptal morphologies and distribution.

Dmp1^Cre^+ cells showed the greatest amount of morphological and positional diversity, likely due to their constitutively active Cre, which would mark a variety of cell types throughout embryonic development. No Dmp1^Cre^+ labeled cells were observed within the smooth muscle, conveniently allowing us to examine the telocytes (green arrows), vascular smooth muscle cells (vSMCs) (blue arrow), and smooth muscle pillars (SMPs) (orange arrow) of the mucosa exclusively (Fig. 2D). These data show that within the small intestine, genetic mouse models frequently mark heterogeneous populations of cells that often overlap with one another. However, such models still can be useful when attempting to isolate or manipulate specific cell types of interest, particularly when faced with a lack of characterized populations or reliable surface markers.

When examining the spatial distribution of our stromal genetic markers, we noticed that several cell types exhibited consistent positioning relative to the intestinal crypts. In most cases, the vasculature primarily traveled in one of two planes: either at the junction of the lamina propria and muscularis, or at the crypt-villus junction (isthmus). This distribution resulted in the cell bodies of Ncad^CreErt2^+ and Nestin^GFP^+ mural cells being stereotypically positioned either at the isthmus, or near the top of the CBC zone close to the *“+4* position” (Fig. 2E-G, Movies S1-3). Telocytes were even more consistent in their distribution, with each crypt generally having 2-3 telocytes at the +4 position and another 2-3 at the isthmus, most often on positioning themselves on opposing sides of the crypt (Fig. 2H). Such regular arrangement would allow for both the selective production of ligands at a given position, and the passive concentration of ligands in specific regions simply due to the geometry of the tissue, as shown by a recent study (Shyer et al., 2015). Our images also revealed that nearly all crypts embed their base within a dense layer of ordered collagen fibers which form the boundary between the mucosa and the muscularis (Fig. S2F). These bundles of collagen fibers are formed by a layer of fibroblasts only a few cells thick, which have recently been described as *Ackr4*+ submucosal fibroblasts and also shown to express key endothelial regulators (Thomson et al., 2018).

### scRNA-Seq of Dmp1^Cre^ populations

Because several recent studies have examined the roles that myofibroblasts and telocytes play in ISC regulation, we also performed scRNA-Seq using the Dmp1^Cre^+ mouse model to better resolve the different contributions these cell types make to the ISC microenvironment. Both Dmp1^Cre^+and Dmp1^Cre^- were included to increase the likelihood of capturing stromal cell types that may have been missed by the genetic models utilized. *Foxl1* and *Pdgfra* were used to identify telocytes (Shoshkes-Carmel et al., 2018), though the reduced sequencing depth of scRNA-Seq techniques likely underrepresents the expression of low-count genes such as transcription factors. Still, most *Foxl1* expression coincided with high levels of *Pdgfra*, identifying the telocyte cluster (Fig. 2I-J). Smooth muscle and myofibroblast cell types were identified using *Acta2*, and *Myh11*. Within the mucosa, Powell et al (2011) describe three primary cell types within the lamina propria that express Acta2: SMPs, which are large bundles of smooth muscle cells running from the muscularis to the tips of the villi and have been suggested to act as ‘pistons’, contracting and pumping fluid out of the lacteals (*Acta2 /Myh11* high); vascular associated smooth muscle cells (vSMCs: *Acta2* high/*Myh11* med); and subepithelial myofibroblasts (SEMFs) (*Acta2/Myh11* low). Interestingly, the designated telocyte cluster also fit the expected SEMF profile, suggesting that these two independently identified populations may either be highly related, or simply different points on a continuum. We also were able to broadly segregate the remaining fibroblastic cells into the pericryptal or villus regions according to a villus specific marker (*Tnc*), or basal mucosa marker (*Postn*) (Fig. 2I) (Bernier-Latmani et al., 2015).

### Wnt and Tgfß/Bmp pathways

We next examined the key stem cell regulatory pathways, focusing on secreted ligands to better understand the signal composition of the mucosal microenvironment. Beginning with the canonical Wnt pathway, the most prevalent ligands associated with canonical signaling were *Wnt2b, Wnt4*, and *Wnt3. Wnt3* showed striking specificities to the epithelial compartment with Paneth and Neurog^Gfp+^ cells producing the majority, in good agreement with previous findings (Sato et al., 2011). The stroma specific Wnts showed notably different patterns between one another, with *Wnt4* being highly specific to the telocytes and villus tips, while *Wnt2b* was expressed sporadically throughout the stroma primarily by fibroblasts (Fig. S3B). *Rspo3* was the highest and most promiscuously expressed R-spondin, with a slight enrichment in the *Ackr4*+ submucosal fibroblasts and high expression levels within the lymphatic vessels (Fig. 3A, S2H, S3B); *Rspo1* also was enriched within the *Ackr4*+ submucosal fibroblasts and a *Thy1*+ enriched subset of the pericryptal stroma, yet was relatively absent from the villus stroma (Fig. 3B, S3B). These data combined with the recent discovery that the various R-spondins can act through different mechanisms suggest that these signals likely play distinct roles under homeostatic conditions but are capable of performing somewhat redundant functions (Park et al., 2018).

**Figure 3.**
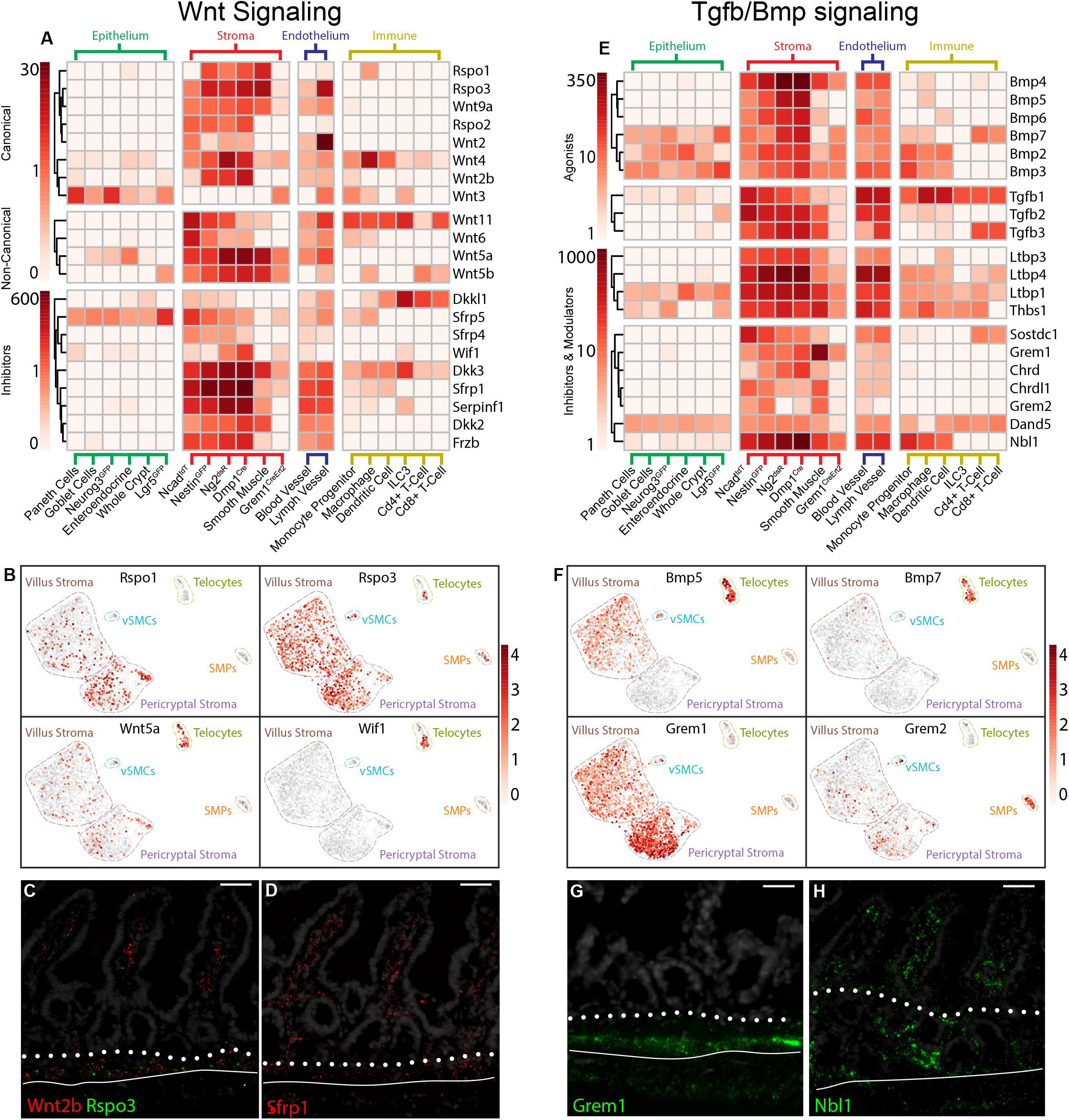
Transcriptome Analysis of the Wnt & Tgfß/Bmp Signaling Pathways. (A) bRNA-Seq analysis of the dominant Wnt signals within the intestinal mucosa reveals that many Wnt signals are highly specific to individual categories. (B) scRNA-Seq analysis of the stroma for key signals within the Wnt pathway. Both spatial and cell type specificity is observed, though redundancies exist among important signals. (C-D) RNAscope in situ detection of *Wnt2b, Rspo3*, and *Sfrp1*. (E) bRNA-Seq analysis of the dominant Tgfb and Bmp signals within the intestinal mucosa. Little categorical specificity was observed. (F) scRNA-Seq analysis of the stroma for key signals of the Bmp pathway. Highly cell type and spatially regulated expression was seen for many members of the superfamily. (G-H) RNAscope in situ detection of Nbl1 and Grem1, the most abundant Tgfb/Bmp inhibitors. Grem1 expression is greatest at the base of the mucosal while Nbl1 expression is highest at the basement membrane of the epithelium. All scale bars are 50μm. See also Fig. S3 for the cognate receptors and scRNAseq results for genes not displayed above.

Of Wnt ligands typically associated with non-canonical pathways, *Wnt5a* and *Wnt5b* were the most abundant. *Wnt5a* was produced primarily by telocytes, while *Wnt5b* showed disperse expression across regions and cell types, somewhat analogous to the R-spondin family (Fig. 3B, S3B). These expression profiles help to explain earlier results showing that ablation of Wnt production from both myofibroblasts and the epithelium is dispensable for ISC function (San Roman et al., 2014), as *Wnt5a* has been shown to be one of the more promiscuously binding Wnt ligands (Dijksterhuis et al., 2015). Interestingly, immune cells produced the vast majority of *Wnt11*, and monocyte progenitors contributed large amounts of *Wnt4*, further adding to the redundancies built into the intestinal niche network (Fig. 3A).

Except for *Sfrp5* in the epithelium and *Dkkl1* in immune cells, the bulk of Wnt inhibitors were produced by the stroma with a slight bias towards the villus, as evidenced by *Sfrp1* and *Dkk2. Dkk2* had a particularly interesting distribution in the scRNA-seq data, with *Dkk2*+ cells being highly concentrated within their cluster (Fig.S3B). Notable exceptions to this spatial bias were the pericryptical *Sfrp4* and the remarkably telocyte-specific *Wif1*. (Fig. 3B,S3B). *Wif1* typically functions as a noncanonical Wnt inhibitor (Vassallo et al., 2016, Skaria et al., 2017), and has been shown to be associated with several cancers and intestinal disorders (Wissmann et al., 2003). This unusually selective expression and clinical relevance make it an attractive candidate for further study. *Sfrp5* was produced by Lgr5^GFP^+ ISCs (Fig. 3A), and is known to be highly expressed by cells at the +4/5 position (Gregorieff et al., 2005). These data suggest it may function as an important regulator of stem cell behavior intrinsic to the epithelium.

Relative to the Wnt pathway, the Tgfß superfamily of ligands had a greater diversity of family members represented and displayed greater specificity in terms of both categorical and cell type expression. *Bmp4* and *Bmp5* were the predominant Bmps within the mucosa, with *Bmp4* showing a slight preference for the villus stroma while *Bmp5* was conspicuously absent from the peri-cryptal stroma, implying stronger spatial than cell type regulation for both (Fig. 3E, S3D). *Bmp3* and *Bmp7* were expressed in both the stroma and epithelium, with telocytes and Lgr5^GFP^+ stem cells being the primary producer for their respective categories; this finding suggests that these ligands are highly epithelial-specific regulators TACs and ISCs (Fig. 3E). The most abundant inhibitors of the Tgfß/Bmp superfamily were *Grem1* and *Nbl1*, with *Chrd* and *Grem2* reaching intermediate levels. In general, these inhibitors were biased towards the pericryptal stroma with the notable exception of *Grem2*, which was primarily expressed by the SMPs (Fig. 3F). Taken together, these data suggest that while there is a general trend of higher Wnt signaling within the pericryptal stroma and higher Bmp signaling within the villus, several localized exceptions exist which have yet to be explored.

### Additional morphogens and growth factors

Telocytes produced the bulk of *Inhba* and *Inhbb*, both of which have been shown to inhibit tumor metastasis and angiogenesis, further implicating telocytes as the first line of defense against epithelial tumor growth and metastasis (Fig. S4D). *Gdf10* and *Gdf6* were unique in their spatial distribution, being the only two members of the superfamily which were both expressed primarily in the pericryptal region, with *Gdf10* expression being practically exclusive to the same *Thy1* enriched population that produces large amounts of *Rspo1* (Fig. 1B, S2H, 4B). In line with previous findings, the epithelium produced most Hedgehog ligands, with *Ihh* constituting the overwhelming majority (Fig. 4A)(Madison et al., 2005). Analogously to other pathways, the highest producer of *Dhh* was the endothelium and their attendant vSMCs, which also produced the greatest volume of the hedgehog inhibitor *Hhip* (Fig. 4A). This ambivalent pattern was repeatedly observed which could imply a mechanism for constraining the signal to a single cell category and may enable the intestine to respond rapidly to abrupt environmental changes.

**Figure 4.**
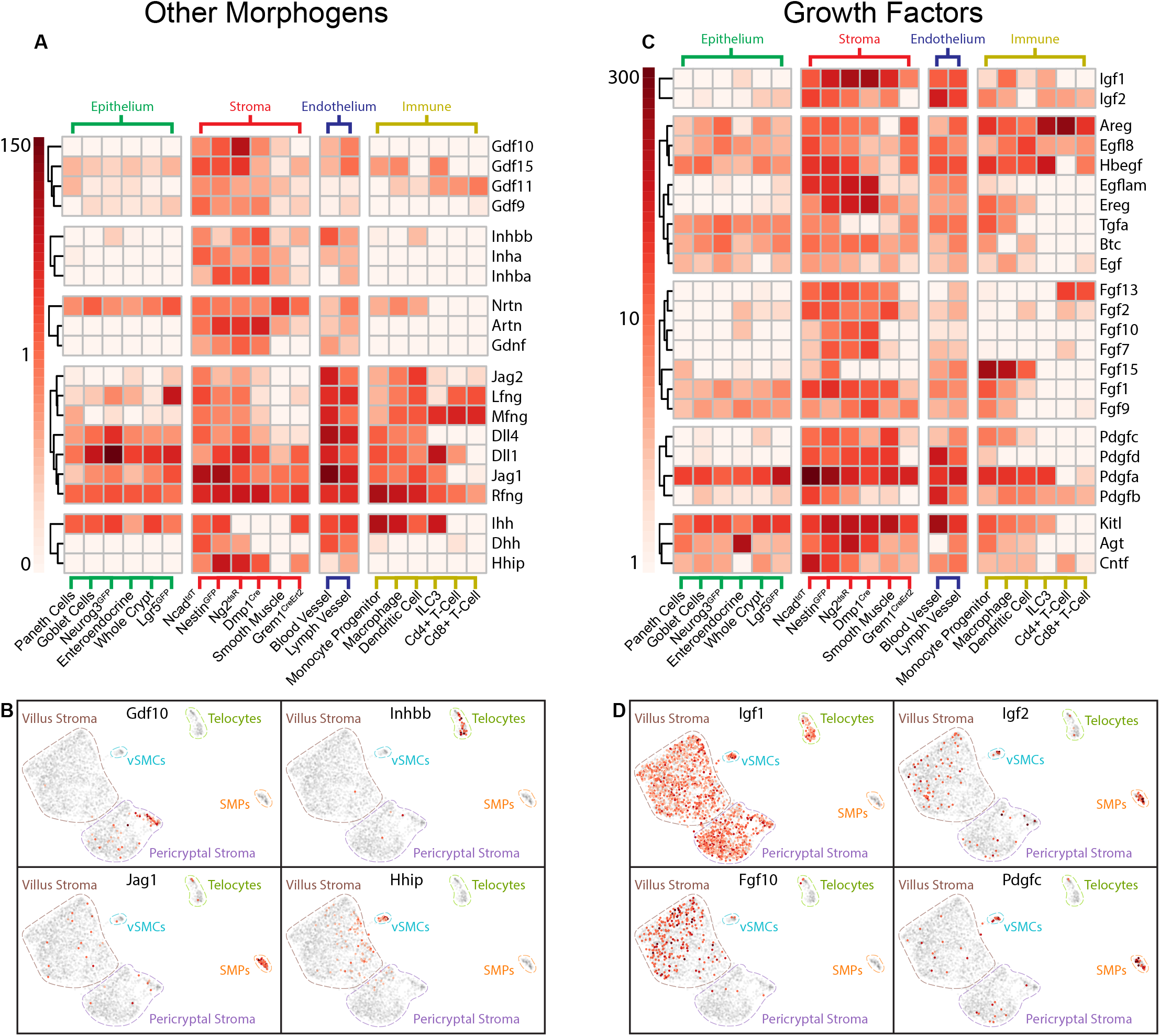
Transcriptome Analysis of Other Morphogens and Growth Factors. (A) bRNA-Seq analysis of additional morphogens expressed at high levels within the intestinal mucosa. (B) scRNA-Seq analysis of important signals reveals a strong cell type specificity for many genes. (C) bRNA-Seq analysis of the predominant growth factors within the intestinal mucosa. Only minor categorical specificity was observed. (D) scRNA-Seq analysis of the predominant growth factors shows strong spatial and cell type specificity. See also Fig. S4 for the cognate receptors and scRNAseq results for genes not displayed above.

Though our bRNA-Seq data suggested that *Igf1* and *Igf2* are widely expressed throughout the stroma and endothelium (Fig. 4C), our scRNA-Seq data revealed inverted expression patterns between the two: virtually all stromal cells produced Igf1, except for the SMPs which instead produced almost all mucosal *Igf2* (Fig. 4D). SMPs likely represent an underappreciated regulatory cell type, as they were the dominant producers of *Hbegf, Pdgfa, Pdgfc* and other growth factors (Fig. 4D, S4D). Of the pathways examined, Egf ligands showed the most epithelial expression, except for *Ereg* and *Egflam* which displayed reciprocal patterns within the stroma (Fig. 4C). Fgf ligand production occurred primarily within the stroma and without obvious specificity, save for *Fgf10* in the villus stroma and *Fgf9* in telocytes (Fig 4C, S4C). Perhaps most interesting was *Fgf15*, which was the most highly expressed Fgf and the only signal observed which was almost exclusively produced by monocytic progenitors (Fig. 4C). SMPs, Ncad^tdT^+, and Lgr5^GFP^+ cells were the highest *Pdgfa* producers, and also happened to be the cells in closest contact to the Pdgfra^High^ telocytes (Fig. 4C, S2A-F). A similar pattern was seen with *Pdgfrb*, with the genetic markers associated with mural cells showing the highest levels of the receptor, while blood endothelial cells produced the most *Pdgfb* and *Pdgfd* (Fig. 4C). This paired Pdgf ligand/receptor expression illustrates just how intimately the structure and molecular profiles of this tissue are intertwined, such that any signaling perturbations will undoubtedly affect multiple cell types.

### Immunomodulatory signals

Surprisingly, chemokines and cytokines were expressed at remarkably high levels and with strong categorical specificities, though little cell type or spatial regulation was apparent (Fig. 5A-B, S5A-B). Epithelially dominant chemokines included *Ccl9, Ccl6*, and to some degree *Ccl25;* stromal specific signals were *Ccl11, Cxcl1, Cxcl12*, and *Cxcl10* (Fig. 5A), with the first two having some spatial preference for the villus (Fig. 5B). *Mif* was expressed at high levels in every cell type examined, and previously has been shown to promote intestinal tumorigenesis, inflammation, and inhibiting macrophage migration (Nobre et al., 2017). *Ccl22, Ccl17, Ccl20, Xcl1* and *Ccl5* were largely immunecell specific, with *Ccl5* being produced in staggering quantities by T-cells; the high level of expression within whole-crypt isolations likely represents T-cells within or adjacent to the crypts (Fig. 5C).

**Figure 5.**
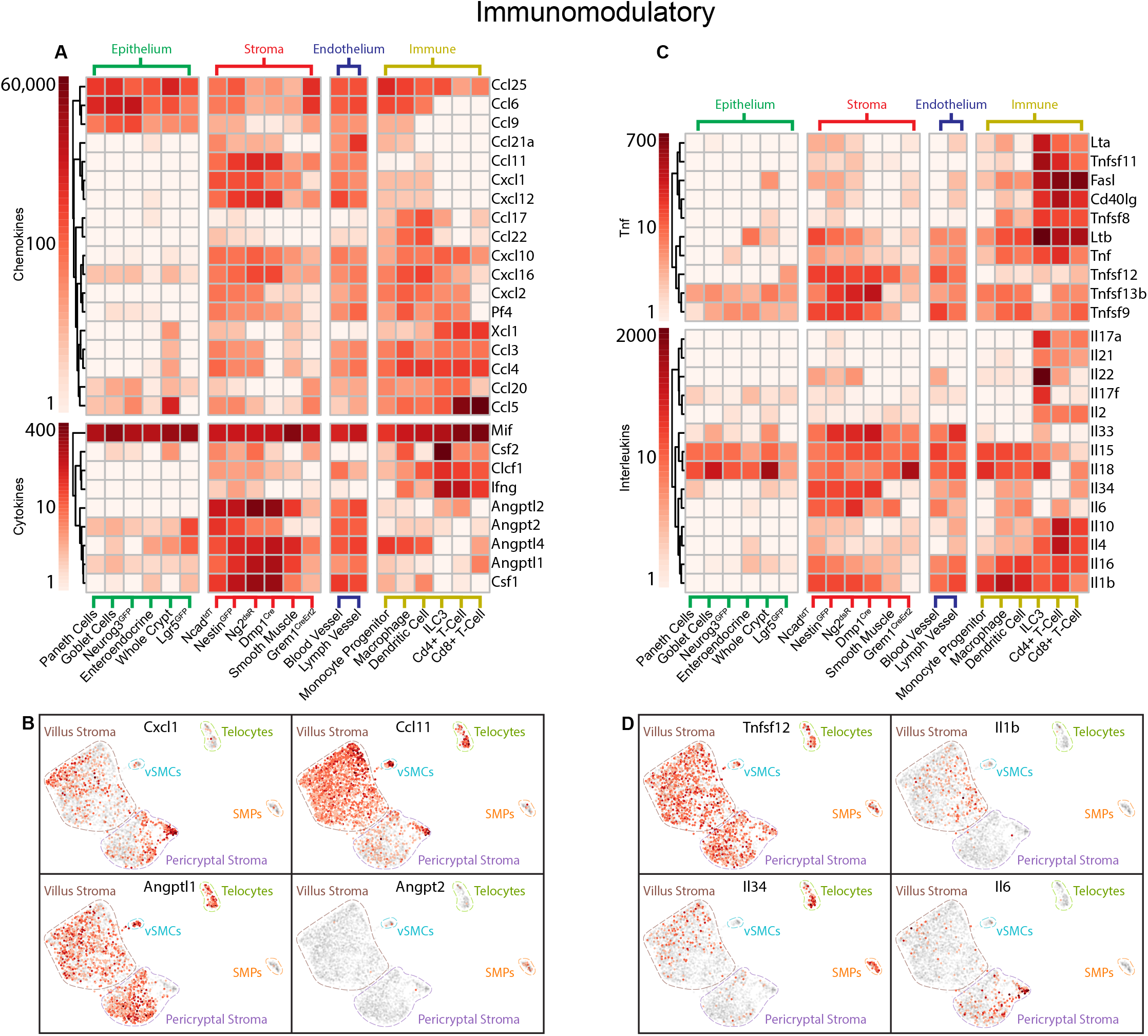
Transcriptome Analysis of Immunomodulatory Pathways. (A) bRNA-Seq analysis of the predominant cytokines and chemokines of the mucosa reveals strong categorical specialization. (B) scRNA-Seq analysis of the stroma. Spatial regulation is present but little cell type specificity was observed. (C) bRNA-Seq analysis of the predominant Tnf and interleukin signals within the mucosa. Though primarily produced by immune cells, some stromal and epithelial expression is observed. (D) scRNA-Seq analysis of important signals shows more spatial than cell type regulation. See also Fig. S5 for the cognate receptors and scRNAseq results for genes not displayed above.

T-cells were the primary producers of both the Tnf and Interleukin ligands. However, a minor stromal preference was observed for *Tnfsf9, Tnfsf12*, and *Tnfsf13b. Il15* and *Il8* were the most heavily and promiscuously expressed interleukins, and *Il18* was notably the only signal observed to reached high levels in the crypts yet was nearly absent from Lgr5^GFP^+ ISCs (Fig. 5C). *Il6, Il33*, and *Il34* were largely stroma-specific, with *Il1b* showing unusual spatial restriction to the villus stroma. A recent study demonstrated that *Il10* promotes ISC self-renewal (Biton et al., 2018), and *Il4s* nearly identical profile suggests it may also play an important role in ISC maintenance (Fig. 5C). ILC3 cells were another major source of interleukins, and they may play an important role in ISC proliferation due in part to the high levels of *Il22* and *Il17* (Withers and Hepworth, 2017).

## Discussion

The diverse functions carried out within the intestinal mucosa require a complicated signaling milieu capable of rapid and proper response to an enormous variety of potential stimuli. However, in order to fully understand how the body responds to these conditions, a solid understanding of the basal homeostatic state must first be established. Our bRNA-Seq analysis results showed that most secreted signaling molecules are expressed in a highly redundant manner across cell types, categories, and regions. Nevertheless, when combined with scRNA-Seq data and immunofluorescent imaging data, an overall organization for the key regulatory signaling components begins to emerge. This more comprehensive view makes it clear that the ISC niche does not consist of a single cell type or molecule, but rather a highly interconnected set of organizational, transcriptional, and morphological characteristics; all of which work collectively to fulfill their supportive and regulatory tasks. The data presented here can help future studies understand how the signaling environment and ISCs themselves respond to specific perturbations within the intestine.

Our imaging results revealed several notable features of stromal cell distribution. Virtually all crypts embed their base in the layer of *Ackr4*+ submucosal fibroblasts which form the basement membrane of the intestinal mucosa (Fig. S2F, Movie S1). These cells produce highly organized bundles of collagen that adopt a serpentine pattern. It is tempting to speculate that the morphology of these fibers provides the cells with the ability to linearize as ingested material stretches the intestine, potentially providing a mechanosensory mechanism for the immediate detection of food passing through that region of the intestine. It is worthwhile to point out the striking morphological, positional, and transcriptional similarity between telocytes originally identified through fluorescent microscopy and the subepithelial myofibroblasts described by Powell et al. (2011). Additionally, our scRNA-Seq data possessed a single cluster which largely fit the description of both SEMFs and telocytes. Though Aoki et al. described telocytes as being negative for *Acta2* and *Myh11*, this claim was supported by only a single immunofluorescent image for each protein (Aoki et al., 2016). One possible explanation for the discrepancy is that Powell et al. (2011) enzymatically enhanced the target signal, while Aoki et al. (2016) used fluorescently labeled secondary antibodies for detection. Expression values spread across several orders of magnitude would cause cells with low expression levels to appear negative via direct detection at concentrations appropriate for the highest levels of expression. However, with enzymatic amplification, positive yet faint staining would develop in *Acta2*^Low^ populations after the *Acta2*^High^ cells had reached saturation. This fact underscores the importance of using identical techniques when determining whether a target of interest is the same as a previously published cell type, or to use additional orthogonal techniques capable of verifying one another.

At similar positions along the crypt-villus axis as the telocytes, a “ring” structure consisting of endothelial and mural cells exists as seen with the Ncad^CreErt2^+ and Nestin^GFP^+ cells. The spatial preferences seen in our genetic models indicate that these two structures may create unique microenvironments. The regular juxtaposition of mural cells and telocytes, along with their high expression levels of the ligand/receptor pairs *Pdgfa* and *Pdgfra*, imply that these cells carry out highly coordinated and cooperative functions that collectively form a “niche network” which can influence ISC function both directly and indirectly (Fig. 4A-B, S2A-F, Movie S2-3). The near exclusive positioning of mucosal Ng2^dsR^+ cells within the villus illustrates that strong compartmentalization of stromal cell types is occurring, though no definitive molecular signatures were seen that may form the mechanistic basis for such organization (Fig. S2B).

Many of the observed structural and organizational characteristics are likely integral to shaping the signaling microenvironment. That nearly all crypts embed themselves within the layer of *Ackr4*+ submucosal fibroblasts suggests that the interactions between these cells are crucial to ISC maintenance and epithelial regeneration. While it has long been known that the presence of Bmp inhibitors like *Grem1, Nog*, and *Chrdl1* in the stem cell zone is essential to ISC self-renewal and maintenance (He et al., 2004, Worthley et al., 2015), the specific cellular source of these signals has remained unknown. Our data support a model in which *Ackr4*+ submucosal fibroblasts are the primary mechanism for shielding ISCs from Bmp signaling through both molecular inhibition and physical occlusion, in addition to enhancing the Wnt signals encountered by the ISCs (Fig. 3B, SB, 6B). Recently it has been demonstrated that Wnt signals spread slowly, primarily via direct cell-cell contact (Farin et al., 2016). This finding, in combination with our structural data, suggest that Paneth cell produced *Wnt3* and the non-canonical ligands from telocytes likely constitute the majority of Wnt ligands encountered by ISCs (Fig. 3A, 6B).

**Figure 6.**
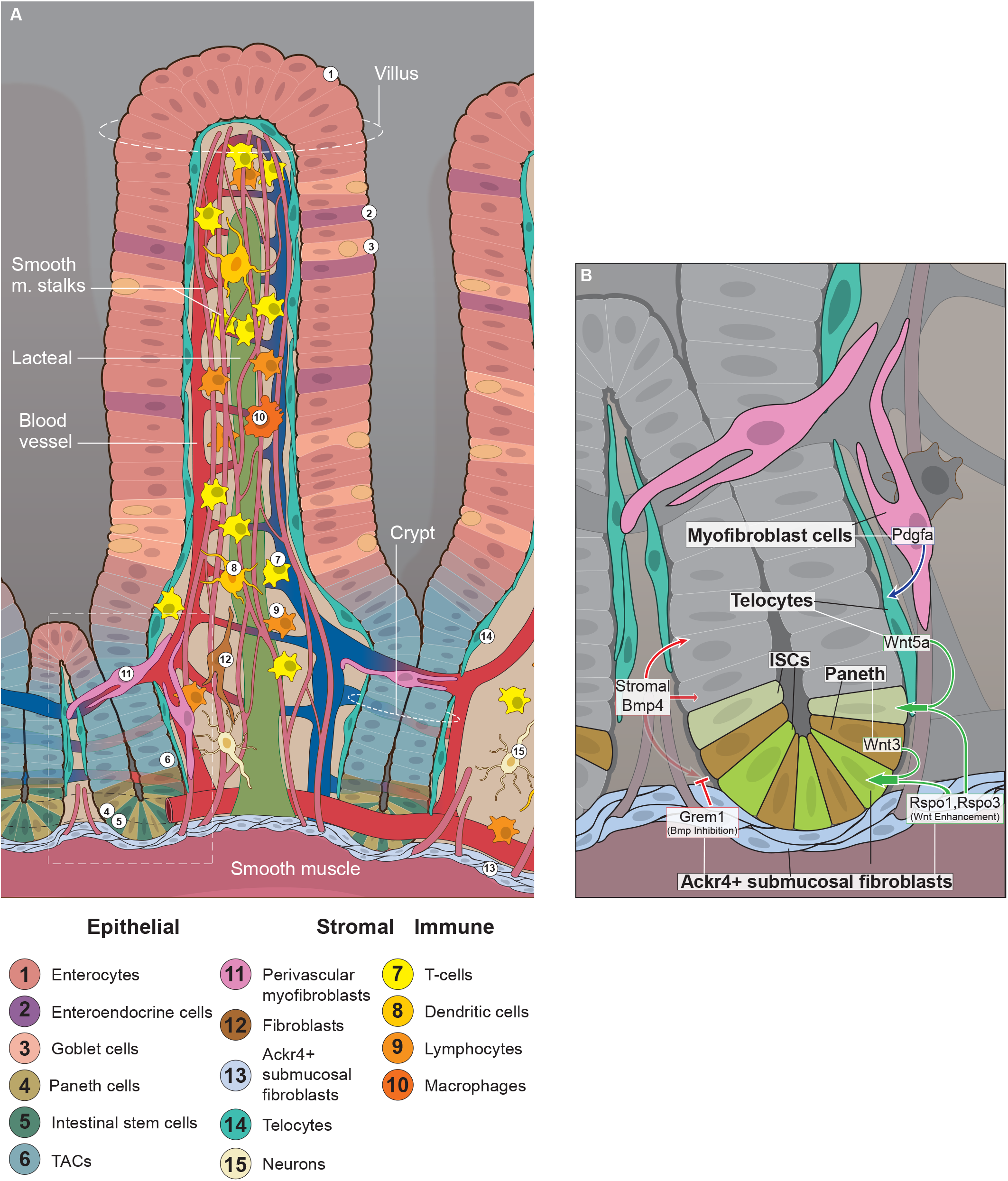
Cellular Organization of the Lamina Propria. (A) Summary of the structural and organizational findings for the Lamina Propria (B) Proposed model for the ISC ‘niche network’ including the key molecular and cellular interactions needed for stem cell regulation.

These observations suggest that the key pathways of Wnt and Bmp, at least for ISCs, are regulated by multiple players which serve distinct roles; telocytes and Paneth cells produce the Wnt ligands needed for self-renewal and expansion, while the *Ackr4*+ submucosal fibroblasts relieve the auto-inhibition of Wnt signals through *Rspo3* to boost pathway activity to levels necessary for self-renewal; meanwhile Bmp pathway activity is reduced through Grem1 to prevent differentiation and inhibit its anti-proliferative effects (Fig. 6B). Such a system has the added benefit of preventing a single aberrant cell from having excessive influence on ISC integrity or inducing carcinogenic transformations. The juxtaposition of telocytes and the observed “ring” structures add yet another layer of regulation and cooperation. Interestingly, Ncad^tdT^+ cells are among the only cells within the intestinal mucosa that produce *Wnt6;* while *Fzd2*, one of the few receptors suggested to interact with *Wnt6* (Dijksterhuis et al., 2015), is the only Fzd receptor widely expressed by telocytes (Fig. 3A, S3A). The juxtaposition of cells expressing highly specific ligand/receptor pairs advises caution when interpreting phenotypes that result from perturbing signaling pathways, as such a tightly orchestrated system is likely to have many indirect effects that could be misinterpreted as direct cell-cell communication.

In general, this receptor-ligand pairing appeared to result in a limited number of autocrine opportunities within the tissue. The endothelial category was the standout exception to this trend, particularly with the lymphatic vasculature. It has been recently noted that the lacteals undergo a constant process of regeneration due to the high amount of mechanical stress and cellular traffic passing through them (Bernier-Latmani et al., 2015). While the authors of that study demonstrated this using *Dll4*, the coexpression of Wnt2/Rspo3/Fzd4 was the only instance of a cell type producing both the cognate ligand and signal enhancer for a receptor it robustly expressed (Fig 3E, S3E). ISCs are not consistently juxtaposed with many lymphatic vessels under normal conditions, yet lymphangiogenesis has been suggested to be an important sequential step in the development of colorectal cancer (CRC). Increasing the exposure of stem cells to the signals produced by the lymphatic vasculature would immediately cause a number of detrimental effects; the oncogenic potential of increased Wnt signaling, the increased ‘self-tolerance’ provided by Ccl21(Kozai et al., 2017), increased inflammation via Il33, and the possible chemo-resistant effects an increase in Tgfb3 could have by reducing proliferative rates of adjacent cells (Fig. 3A&E, 5C) 4. It is therefore interesting to note that while we could not identify any clear roles the endothelium might play within the homeostatic ISC niche, the potential oncogenic and autoimmune effects from their involvement are of obvious concern.

Another potentially underappreciated element is the geometry of the tissue, as it has been shown that the shape of a structure can concentrate or dilute ligands without any change in expression levels (Shyer et al., 2015). We expect that the pericryptal enrichment of stromal signals, in combination with epithelial contributions like *Sfrp5*, inhibits cell proliferation and support relatively quiescent stem/progenitor cells within the +4 region (Fig. 3A)(Powell et al., 2012, Gregorieff et al., 2005). As the TACs move upwards past the first telocyte, mural, and endothelial cell “ring”, they encounter increasing levels of other growth factors, causing them to undergo rapid proliferation and begin differentiating. At the isthmus, the second ring structure can provide additional instructive signals, inducing TACs to exit the cell cycle and complete the differentiation process.

We also found that ILC3 cells express the highest *Il22* and *Lif*, both of which can support ISCs’ survival and proliferation (Fig 5C)(Lindemans et al., 2015). Intriguingly, we also show that ILC3 cells express high levels of *Il17a* and *Il17b*, suggesting a relationship between ILC3 cells and Th17 cells (Chung et al., 2006). Macrophages have been reported to support crypt cells (Pull et al., 2005). Our group also has observed that the distribution of macrophages is different under homeostatic and stressed conditions; Normally CD68^+^ macrophages localize to the villus region, however upon irradiation they migrate towards the bottom of the crypts. This migration may allow them to promote ISC function and regeneration of the intestinal epithelial cells following damage or disease. We have additionally verified some of the key signals using RNAscope including *Rspo3, Grem1*, and *Wnt2b*, but many other signals need to be verified for their cellular and regional distribution in the future (Fig. 3C,D,G,H). This study, therefore, mainly serves as a resource by providing transcriptomic data with different types of niche throughout the intestinal mucosa.

Though each of these pathways is organized differently, several common features emerge. First, while ligand expression values varied greatly between the various pathways, expressed ligands within a given family were generally expressed within the same dynamic range. In addition, secreted inhibitors and modulators for a given pathway were expressed at substantially higher levels, generally an order of magnitude greater at least. Second, highly cell-type specific production of ligands was less common than categorical specificity; genes highly expressed by one category were unlikely to be expressed at similarly high levels in another category. Unfortunately, of the genes that were highly specific to a single cell type, it was unlikely for a single genetic or surface marker to overlap with a given ligand. This highlights the importance of using scRNA-Seq to determine whether changes are concentrated within a subset of cells, or evenly distributed across the population. Another interesting trend observed was the frequent ambivalent expression of both agonist and antagonists within a single cell type. For example, telocytes were among the highest expressors of both Wnt ligands and inhibitors, and Nestin^GFP^+ cells presenting a similar pattern with the hedgehog pathway.

In conclusion, this study provided a holistic evaluation of the predominant cell types and signals present within the intestinal mucosa, above and beyond the role of any single cell type or signal. By utilizing bRNA-Seq to profile 20 different mucosal cell types covering four major cell categories (epithelial, stromal, endothelial and immune) and examining the stromal signaling environment via scRNA-Seq, we have been able to provide a more comprehensive view of the signaling microenvironment within the intestinal mucosa, identifying primary signals for the major ISC regulatory pathways and their respective cellular sources. Our analysis leads to the conclusion that a ‘niche network’ exists, in that no single cell type is responsible for ISC self-renewal or expansion. Rather our findings demonstrate that each cell type within the network carries out its own specific functions, in a highly cooperative and coordinated manner (Fig. 6).

## Acknowledgements

Research reported in this publication was supported by National Institute of Diabetes and Digestive and Kidney Disorders (NIDDK) and National Institute of Allergy and Infectious Diseases (NIAID) of the National Institutes of Health under grants number U01DK085507, DK085547, DK085532, DK085535, DK085527, DK085508, DK085551, DK085525, DK085551, and DK085527. The content is solely the responsibility of the authors and does not necessarily represent the official views of the National Institutes of Health.

## Author contributions

Conceptualization, D.M.H., M.W., S.H.,M.M. and L.L.; Methodology, D.M.H., X.H., L.L., S.C. and A.P.; Formal Analysis, D.M.H. and S.C.; Investigation, D.M.H., P.I.M.G., Y.C., D.H., D.T., and X.H.; Writing – Original Draft, D.M.H. & L.L.; Writing – Review & Editing, D.M.H., B.J.O., J.C.N., M.W., T.C.W and L.L; Resources, M.G.M, T.C.W, M.W., C.K., C.M.D, C.W.H., J.Y., J.L., and A.M.H.; Data Curation, S.C., A.P.; Visualization, D.M.H.; Supervision, L.L. X.H.; Project Administration, M.G.M., B.J.O., and J.C.N.; Funding Acquisition, L.L. via ISCC and SIMR.

## STAR Methods

### Key Resources Table

### Lead Contact and Materials Availability

Further information and requests for reagents should be directed to and will be fulfilled by the Lead Contact, Linheng Li.

### Experimental Model and Subject Details

#### Mice

The mice used in this study of the following strains were housed in the animal facility at the Stowers Institute for Medical Research (SIMR) and were handled according to SIMR and National Institutes of Health (NIH) guidelines. All procedures were approved by the Institutional Animal Care and Use Committee (IACUC) of the SIMR. Tg(Cspg4-DsRed.T1)1Akik/J, mice were obtained from The Jackson Laboratory. N-cad-CreERT, and N-cad-TdT mice were generated by Applied StemCell, Inc. The Nestin-GFP transgenic mouse line was previously described (Mignone et al., 2004). The B6N.FVB-Tg(Dmp1-cre)1Jqfe/BwdJ mice were a generous gift from the lab of Dr. Sarah Dallas. To induce N-cad-CreERT; R26-tdT mouse, TMX (Sigma) was injected intraperitoneally at 2mg per injection for 3 days.

Cells from the B6.Cg-Tg(Grem1-cre/ERT)3Tcw/J strain were provided by the lab of Dr. Timothy Wang.

### Method Details

#### Whole mount Immunofluorescence

Whole mount samples were prepared, stained, and imaged by adapting the protocols established by the labs of T. Petrova and I. Nathke (Appleton et al., 2009, Bernier-Latmani and Petrova, 2016). Briefly, extracted intestines were fixed in either 15% picric acid, 0.5% PFA, and 0.1 M sodium phosphate, or in 4% PFA in PBS overnight at 4°C. Samples were then washed and cryoprotected in 20% sucrose and 10% glycerol. Pieces roughly 1 cm^2^ were blocked and permeabilized in buffer that contained 0.3% BSA, 5% donkey serum, and 0.1% Triton X-100. Primary antibodies were incubated at 4°C for 18-48 hours, and secondary antibodies were incubated at 4°C overnight. Tissue was cleared using 2,2’-Thiodiethanol and cut into strips 1-2 villi thick, then mounted in 97% TDE 3% PBS. Images were acquired using either a Zeiss LSM-780 DS or Nikon 3P microscope and analyzed using FIJI (Schneider et al., 2012, Schindelin et al., 2012). In the case of two-photon microscopy, we followed a previously developed technique (Appleton et al., 2009).

#### Flow Cytometry

##### Stromal, endothelial, and immune cell isolation

For both bRNA-Seq and scRNA-Seq for the stromal, endothelial, and immune categories, cells were isolated using a modified version of the protocol established by the Lucie Peduto lab (Stzepourginski et al., 2015). The intestine was extracted, linearized and washed with HBSS without calcium or magnesium. The smooth muscle was then removed by placing the intestine villi-down onto a plate lined with PDMS, then using a razor blade to peel the muscle away from the mucosa. The epithelium was removed by placing the mucosa into HBSS without calcium or magnesium + 10mM EDTA, then placed on a rocker for 15 minutes at room temperature; samples were then shaken for 1-2 minutes and the supernatant discarded until a clear supernatant was achieved (typically 2-3 repeats). The lamina propria was then cut into as small of pieces as possible and placed in 10mLs of digestion buffer consisting of DMEM with Liberase TL at 0.5 Wunsch units and DNaseI at 2 units/mL for 20 minutes at 37° under gentle agitation. After 20 minutes, the supernatant was collected and moved to a quenching tube, and a fresh 5 mls of dissociation buffer was added to the remaining tissue and the process repeated once. The digestion was quenched by adding one volume of DMEM with 10% FBS and 1 unit/ml of DNaseI and placing on ice. The single-cell suspension was then strained using a 70μM filter, pelleted, and resuspended in staining buffer consisting of HBSS + 3% FBS at 10e6 FSC/SSC gate events/mL. Single cell suspensions were stained for 30 minutes on ice then washed and sorted.

##### Epithelial cell isolation

Epithelial cells were isolated as previously described (Wang et al., 2013). Briefly, intestines were extracted, linearized and washed with HBSS without calcium or magnesium. Samples were then cut into small pieces, placed into HBSS without calcium or magnesium with 0.5M EDTA, and incubated in a 50ml conical tube on ice for 20 minutes. Afterwards, Tubes were shaken for 5 minutes and the supernatant run through a 70μm strainer. The resulting suspension was spun down, the supernatant removed, and the pellet resuspended in TryPLE Express containing 0.5 mM NAC. The pellet was resuspended by pipetting, and samples were incubated at 37°C for 5 minutes with gentle shaking and pipetting to aid dissociation. Samples were then transferred to a tube containing ice cold ADMEM/F12 medium for washing and shaken to discourage aggregation. After one minute, tubes were spun at 600g for 5 minutes at 4°C, and the pellet was resuspended in ice cold HBSS with 3% FBS and stained for 30 minutes on ice then washed and sorted.

#### bRNA-seq RNA extraction and cDNA preparation

mRNA-Seq libraries were generated from approximately 1ng of total RNA, as assessed using the Bioanalyzer (Agilent, Santa Clara, CA). cDNA and libraries were made according to the manufacturer’s directions for the SMART-Seq v4 Ultra Low Input RNA Kit (Takara Bio, Kusatsu, Shiga Prefecture, Japan, Cat. No. 634890) and Nextera XT DNA Library Prep Kit (Illumina, Cat. No. FC-131-1096). Resulting short fragment libraries were checked for quality and quantity using the Bioanalyzer (Agilent) and Qubit Fluorometer (Life Technologies). Libraries were pooled, requantified and sequenced as 50bp single reads on 2 lanes of a high-output flowcell using the Illumina HiSeq 2500 instrument.

#### Single cell capture and cDNA preparation

Dissociated cells, having been sorted in phosphate-buffered saline with 0.04% BSA, were assessed for concentration and viability via a Nexelom Cellometer Auto T4. Cells, having been deemed to be at least 50% viable, were loaded on a Chromium Single Cell Controller (10x Genomics, Pleasanton, CA), based on live cell concentration, with a target of 2,000-3,000 cells per sample. Libraries were prepared using the Chromium Single Cell 3’ Library & Gel Bead Kit v2 (10x Genomics) according to manufacturer’s directions. Resulting short fragment libraries were checked for quality and quantity using an Agilent 2100 Bioanalyzer and Invitrogen Qubit Fluorometer. Libraries were pooled at equal molar concentrations and sequenced to a depth necessary to achieve 40,000-55,000 mean reads per cell - ~125M reads each - on an Illumina HiSeq 2500 instrument using Rapid SBS v2 chemistry with the following paired read lengths: 26 bp Read1, 8 bp I7 Index and 98 bp Read2.

#### In situ

Extracted intestines were cryoprotected in 30% sucrose overnight and embedded in optimum cutting temperature (OCT) compound (Tissue Tek) then sectioned 10μm thick. Single-molecule in situ hybridization was performed using Advanced Cell Diagnostics RNAscope^®^ Multiplex Fluorescent Reagent Kit v2 (Cat. No. 323100). The ISH protocol was followed according to the manufacturer’s instructions.

### Quantification and Statistical Analysis

#### QC, alignment, and counting bRNA-seq data

Illumina Primary Analysis version RTA 1.18.64 and bcl2fastq2 v2.18 were run to demultiplex reads for all libraries and generate FASTQ files. Raw reads were demultiplexed into Fastq format allowing up to one mismatch using Illumina bcl2fastq2 v2.18. Reads were aligned to UCSC genome mm10 with TopHat v2.0.13, default parameters. FPKM values were generated using Cufflinks v2.2.1 with “-u –max-bundlefrags 100000000.” Read counts were generated using HTSeq-count with “-m intersection-nonempty.” Three to four replicates were sequenced for each population.

#### Analysis of bRNA-seq data

#### QC, alignment, and counting scRNA-seq data

cDNA libraries were sequenced as paired-end reads on Illumina HiSeq 2500 machine. Raw sequencing data were processed using 10X Genomics software cellranger. Reads were demultiplexed from Bcl into Fastq file format using cellranger mkfastq. Genome index was built by cellranger mkref using mouse genome mm10, ensembl 87 gene model. Data were aligned by STAR aligner and cell counts tables were generated using cellranger count function with default parameters.

#### Normalization and analysis of scRNA-seq data

The R package Seurat was used for quality control, nonlinear dimensionality reduction and clustering. Dmp1^Cre^+ and Dmp1^Cre^- datasets were used to create Seurat objects which were then combined using the MergeSeurat function. Genes expressed in < 3 cells were dropped, while only cells having a unique gene count between 1000 and 3000 with less than 15% mitochondrial gene content were included in the analysis.

NormalizeData was used to log normalize and multiplied by a factor of 10^4^. The variable genes used in dimensional reduction and clustering were determined using FindVariableGenes function, with an x.low.cutoff of 0.1 and an x.high.cutoff of 1. To reduce the influence of Immediate early genes (IEGs) on clustering, the top 15 IEGS represented in the data were used in the AddModuleScore function to score each cell based on their IEG content. Data was scaled linearly and the effects of IEG content, mitochondrial gene percentage, and UMI count were regressed out at the ScaleData step using the vars.to.regress parameter. Principal Component Analysis was chosen for dimensional reduction, and the first 20 principle components were used for the remaining analysis based on the results of the Jack Straw plot results. Clusters were identified using the FindClusters function and plotted using the Uniform Manifold Approximation and Projection (UMAP) dimensional reduction method. The ability of each gene to identify a cluster was determined using the ROC test in the FindAllMarkers function; genes were given a score between −1 (perfect negative correlation with a cluster) and 1 (perfect correlation with a cluster), and genes with both a power above 0.1 (min.pct) and log fold change of greater than 0.25 (logfc.threshold) were chosen.

### Data and Code Availability

#### Data Resources

The data can be retrieved from GEO database with accession number GSE139095.

## Supplemental Figure Titles and Legends

**Figure S1. Table of Isolated Cell Types and Their Corresponding Functional genes**

(A) Table showing the list of cells used for the bRNA-seq results and the markers used in FACS isolation.

(B) Heatmap of Functional genes demonstrating the expected populations were obtained in Figure 1

**Figure S2. Colocalization, Pericryptal organization, and ID genes for stromal populations in Figure 2**

(A) Whole mount composite view of Fig. 1A-B, representing Nestin-GFP;Ncad-Cre;R26R-tdTomato single and double positive cells.

(B) Whole mount view of Nestin-GFP;Ng2-dsRed single and double positive cells.

(C) Morphology and distribution of Ncad-Cre marked cells within the pericryptal region.

(D) Morphology and distribution of Nestin-Gfp marked cells within the pericryptal region. Asterisk marks location where a thin projection from a Nestin-GFP+ cell is encompassed by a larger Ncad-Cre+ tube-like structure.

(E) Morphology and distribution of Nestin-Gfp;Ncad-Cre;R26R-tdTomato marked cells within the pericryptal region.

(F) Orthogonal view of Dmp1-Cre;R26R-tdTomato double-positive cells and second harmonic generation (SHG) signals within the pericryptal region. Z1-4 are slices that correspond to their respective dashed line in the first panel; the dotted line in the z frames represent the location of the top frame.

Median filter has been applied to panels A-D to reduce noise, and gamma has been adjusted on panels B-D to visualize both large and small structures within the same image. All scale bars are 50μm.

**Figure S3. scRNAseq of remaining bRNAseq genes and Receptor Expression of Wnt & Bmp Pathways in Figure 3**

(A-B) Receptors for the corresponding ligands in Figure 3.

(C-D) scRNAseq results for remaining genes in Figure 3.

**Figure S4. scRNAseq of remaining bRNAseq Genes and Receptor Expression for Additional Morphogens & Growth Factors in Figure 4**

(A-B) Receptors for the corresponding ligands in Figure 4.

(C-D) scRNAseq results for remaining genes in Figure 4.

**Figure S5. scRNAseq of Remaining bRNAseq Genes and Receptor Expression For Immunomodulatory Signals in Figure 5**

(A-B) Receptors for the corresponding ligands in Figure 5.

(C-D) scRNAseq results for remaining genes in Figure 5.

## Supplemental Video Titles and Legends

**Movie S1. Structure and Distribution of Nestin-GFP+ cells and Ackr4+ fibroblasts**

Nestin cells are green and SHG signal is in blue. At 0:05 a clipping plane removes the basement membrane of the lamina propria to reveal the basement membrane at the stroma-epithelial junction. The upper and lower planes of mural cells can be seen beginning at 0:17.

**Movie S2. Structure and Distribution of Ncad^Cre^+ cells**

The magnified view demonstrates the way these cells encircle the endothelium (large tubes) and telopode-like projections from Nestin-GFP+ cells (thinner tubes)

**Movie S3. Spatial relationship between Ncad^Cre^+ cells and Lgr5+ ISCs**

Ncad^Cre^+ (red) and Lgr5^DTR-GFP^+ (green) are in close proximity to one another, with Ncad cells adopting a stereotypical position at the isthmus and muscularis-mucosal junction.

