## Supplemental figures for "A Holistic Analysis of the Intestinal Stem Cell Niche Network"

**A**

| Compartment | Name Used | Positive markers | Negative markers |
|---|---|---|---|
| Epithelial | Paneth Cells | UEA-1;CD24(lo) | |
| | Goblet Cells | UEA-1 | CD24 |
| | Neurog3 | Genetic marker | |
| | Enteroendocrine | UEA-1;CD23 | |
| | Whole Crypt | Mechanically isolated | |
| | Lgr5 | Lgr5-GFP | |
| Stromal | Ncad | Cdh2-CreERT;R26-LSL-TdTomato | CD31;CD45;CD326;TER-119 |
| | Nestin | Nes-GFP | CD31;CD45;CD326;TER-119 |
| | Ng2 | Cspg4-DsRed | CD31;CD45;CD326;TER-119 |
| | Dmp1 | Dmp1-Cre;R26-LSL-TdTomato | CD31;CD45;CD326;TER-119 |
| | Smooth Muscle | Genetic marker | |
| | Grem1 | Grem1-CreERT;R26-LSL-TdTomato | CD31;CD45;TER-119 |
| Endothelial | Blood Vessels | CD31 | CD45;CD326;TER-119 |
| | Lymph Vessels | CD31;Pdpn | CD45;CD326;TER-119 |
| Immune | Macrophages | CD11b;CD45 | CD31;CD326;TER-119;CD11c |
| | Dendritic Progenitor | CD11b;CD11c;CD45 | CD31;CD326;TER-119 |
| | Dendritic Cells | CD11c;CD45 | CD31;CD326;TER-119;CD11b |
| | Innate Lymphoid 3 | RORyt;CD45 | CD11b;CD11c;B220;CD3 |
| | Cd4+ T-Cells | CD3;CD4 | CD326 |
| | Cd8 T-Cells | CD3;CD8 | CD326 |

**B**

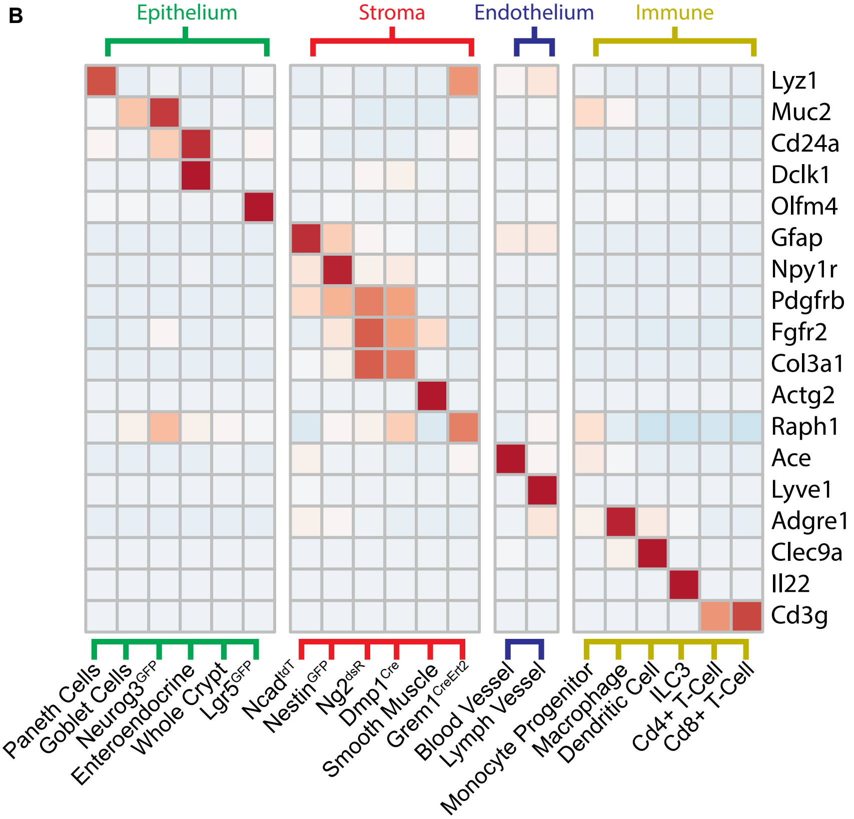

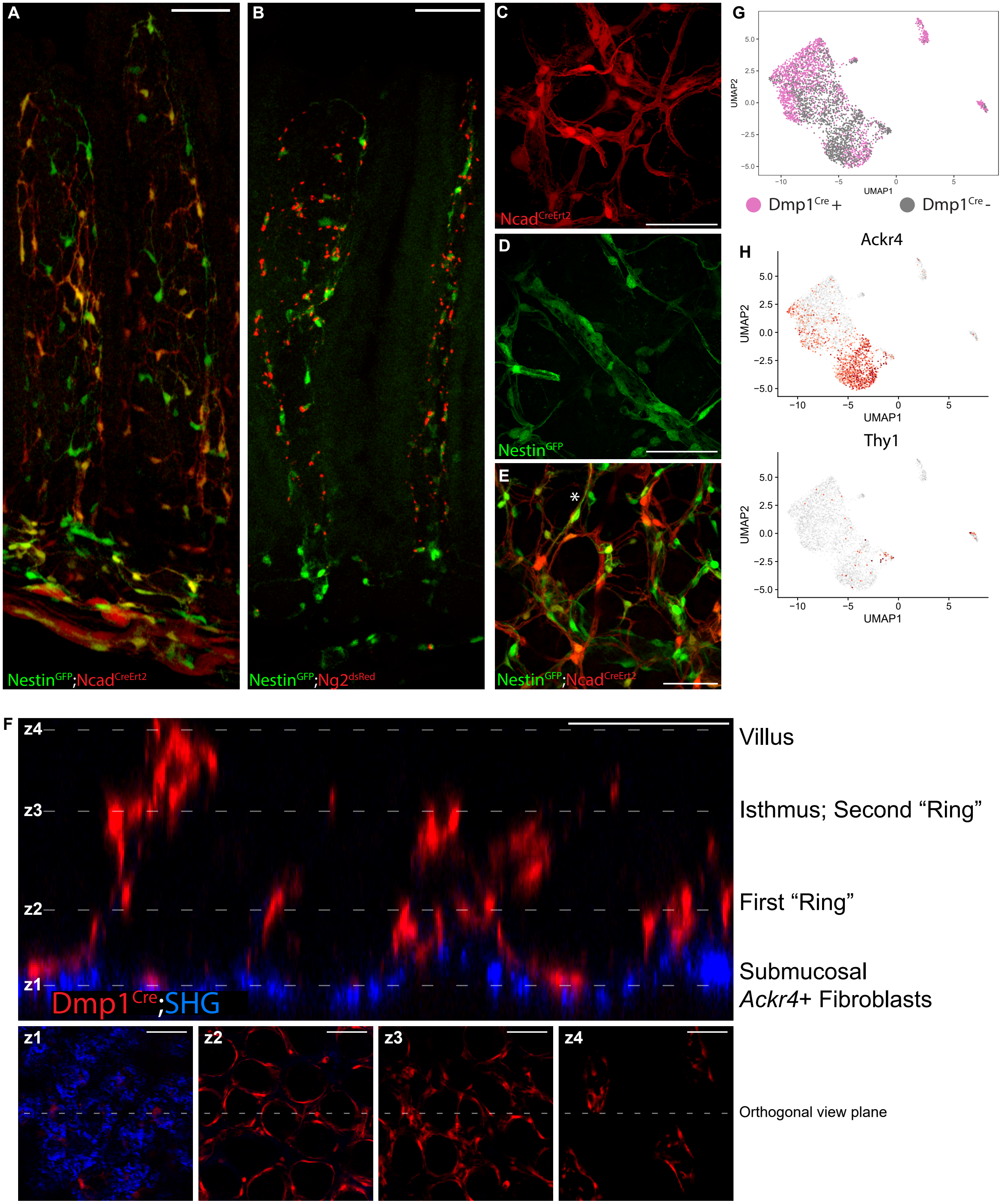

A
B
C
D
E
F
G
H
Nestin$^{GFP}$;Ncad$^{CreErt2}$
Nestin$^{GFP}$;Ng2$^{dsRed}$
Ncad$^{CreErt2}$
Nestin$^{GFP}$
Nestin$^{GFP}$;Ncad$^{CreErt2}$
UMAP2
UMAP1
Dmp1$^{Cre}$+
Dmp1$^{Cre}$-
Ackr4
Thy1
z4
z3
z2
z1
Dmp1$^{Cre}$;SHG
Villus
Isthmus; Second “Ring”
First “Ring”
Submucosal
*Ackr4*+ Fibroblasts
Orthogonal view plane

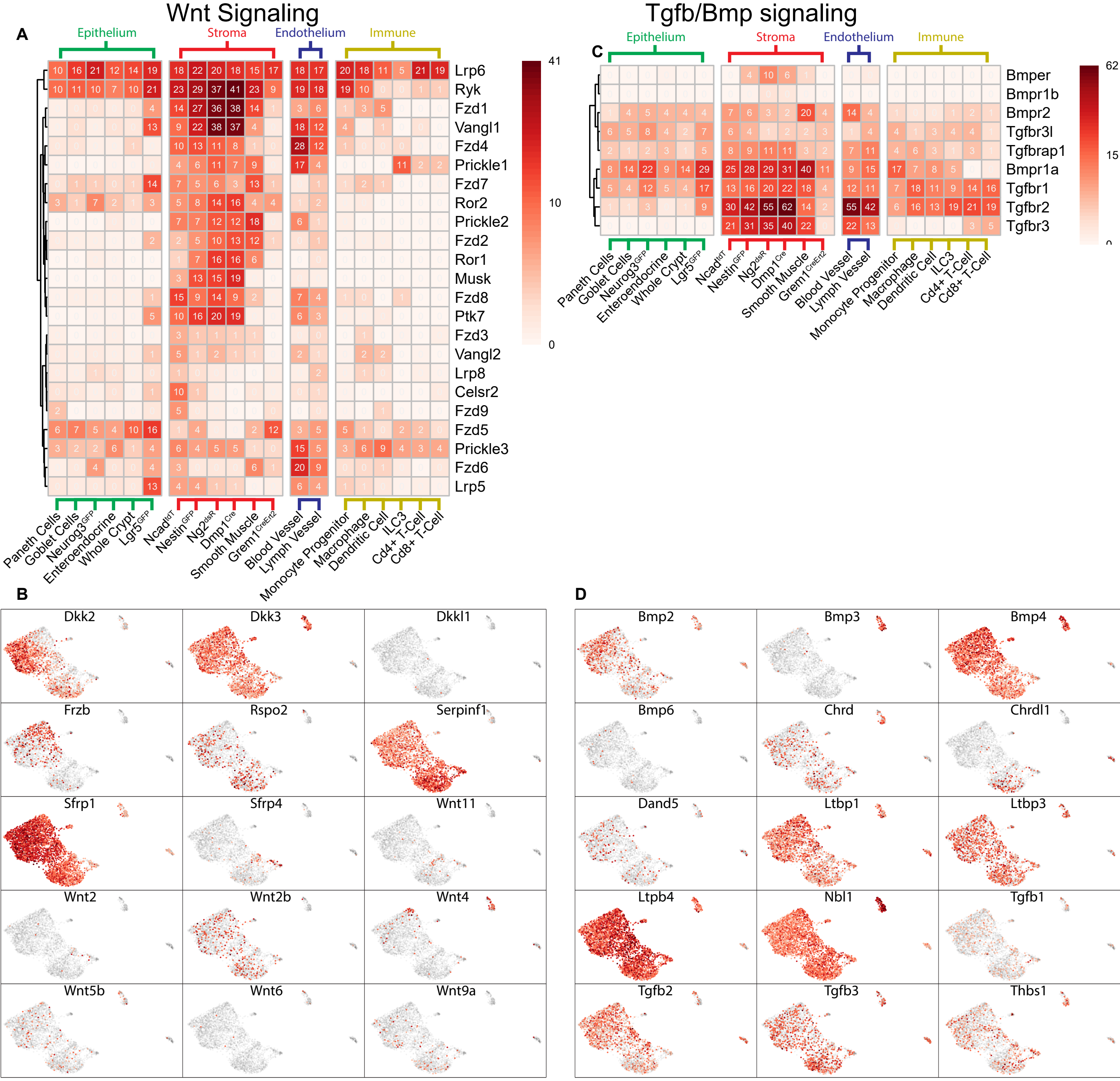
Wnt Signaling
A
Epithelium
Stroma
Endothelium
Immune
Lrp6
Ryk
Fzd1
Vangl1
Fzd4
Prickle1
Fzd7
Ror2
Prickle2
Fzd2
Ror1
Musk
Fzd8
Ptk7
Fzd3
Vangl2
Lrp8
Celsr2
Fzd9
Fzd5
Prickle3
Fzd6
Lrp5
41
10
0
Paneth Cells
Goblet Cells
Neurog3GFP
Enteroendocrine
Whole Crypt
Lgr5GFP
NcadtdT
NestinGFP
Ng2dsR
Dmp1Cre
Smooth Muscle
Grem1CreErt2
Blood Vessel
Lymph Vessel
Monocyte Progenitor
Macrophage
Dendritic Cell
ILC3
Cd4+ T-Cell
Cd8+ T-Cell
Tgfb/Bmp signaling
C
Bmper
Bmpr1b
Bmpr2
Tgfbr3l
Tgfbrap1
Bmpr1a
Tgfbr1
Tgfbr2
Tgfbr3
62
15
0
B
Dkk2
Dkk3
Dkkl1
Frzb
Rspo2
Serpinf1
Sfrp1
Sfrp4
Wnt11
Wnt2
Wnt2b
Wnt4
Wnt5b
Wnt6
Wnt9a
D
Bmp2
Bmp3
Bmp4
Bmp6
Chrd
Chrdl1
Dand5
Ltbp1
Ltbp3
Ltpb4
Nbl1
Tgfb1
Tgfb2
Tgfb3
Thbs1

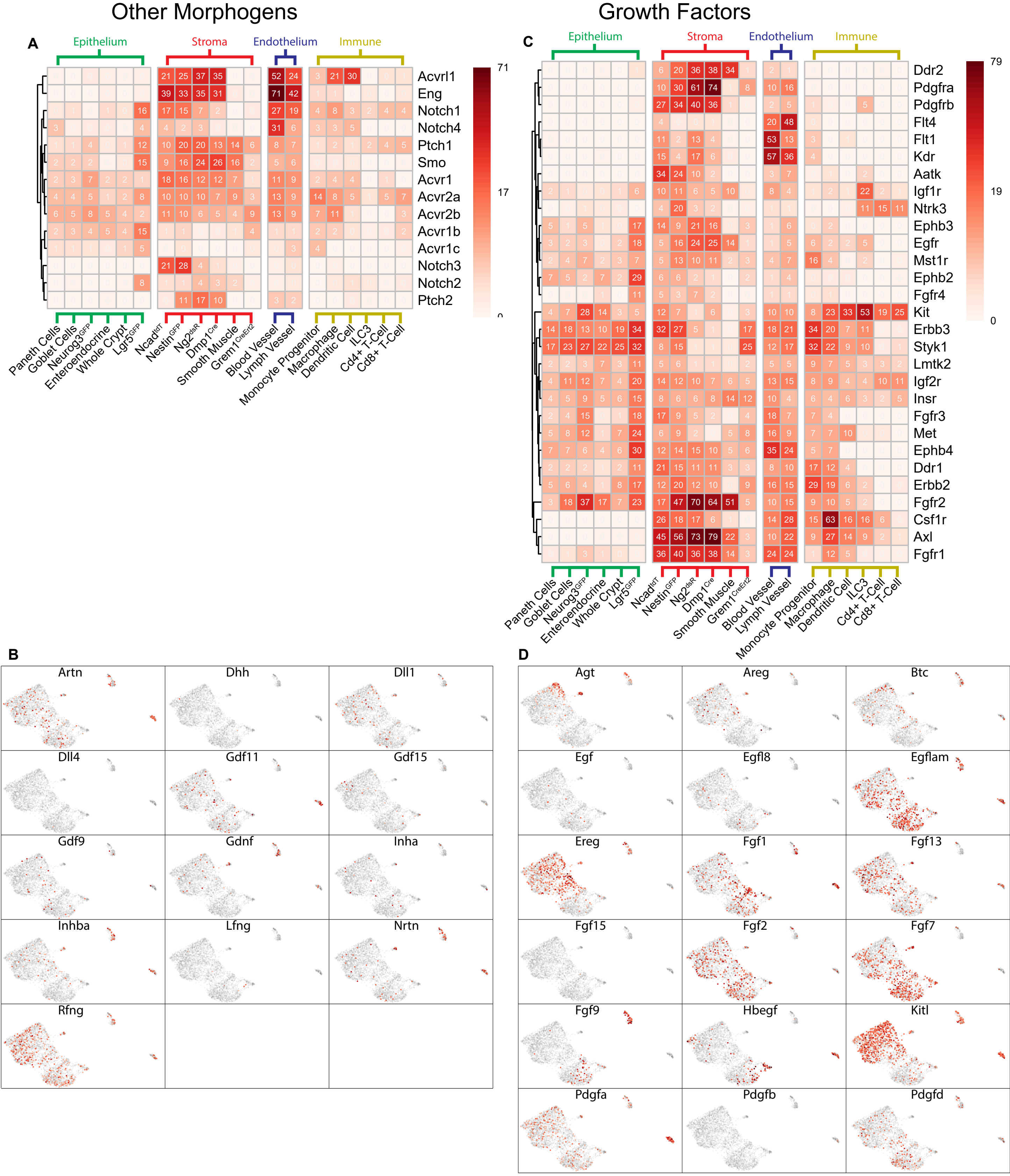

Other Morphogens
Growth Factors
A
B
C
D
Epithelium
Stroma
Endothelium
Immune
Acvrl1
Eng
Notch1
Notch4
Ptch1
Smo
Acvr1
Acvr2a
Acvr2b
Acvr1b
Acvr1c
Notch3
Notch2
Ptch2
71
17
0
Paneth Cells
Goblet Cells
Neurog3GFP
Enteroendocrine
Whole Crypt
Lgr5GFP
NcadtdT
NestinGFP
Ng2dsR
Dmp1Cre
Smooth Muscle
Grem1CreErt2
Blood Vessel
Lymph Vessel
Monocyte Progenitor
Macrophage
Dendritic Cell
ILC3
Cd4+ T-Cell
Cd8+ T-Cell
Ddr2
Pdgfra
Pdgfrb
Flt4
Flt1
Kdr
Aatk
Igf1r
Ntrk3
Ephb3
Egfr
Mst1r
Ephb2
Fgfr4
Kit
Erbb3
Styk1
Lmtk2
Igf2r
Insr
Fgfr3
Met
Ephb4
Ddr1
Erbb2
Fgfr2
Csf1r
Axl
Fgfr1
79
19
0
Artn
Dhh
Dll1
Dll4
Gdf11
Gdf15
Gdf9
Gdnf
Inha
Inhba
Lfng
Nrtn
Rfng
Agt
Areg
Btc
Egf
Egfl8
Egflam
Ereg
Fgf1
Fgf13
Fgf15
Fgf2
Fgf7
Fgf9
Hbegf
Kitl
Pdgfa
Pdgfb
Pdgfd

### Immunomodulatory

A

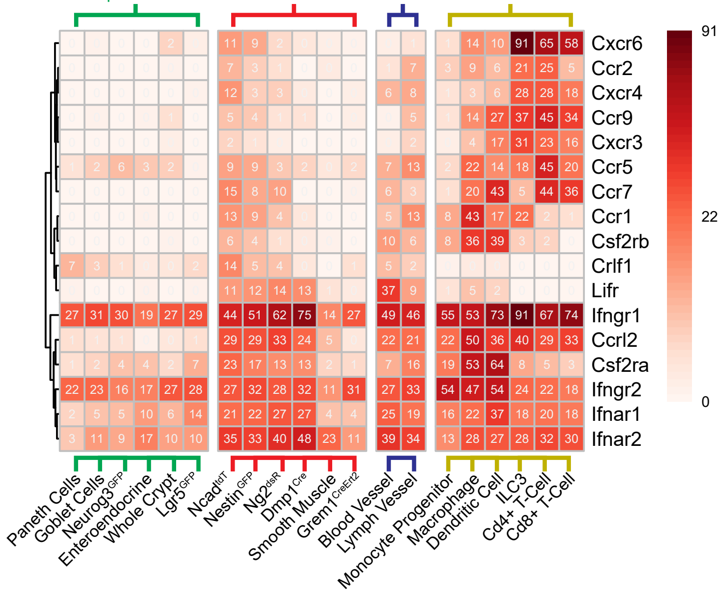


D

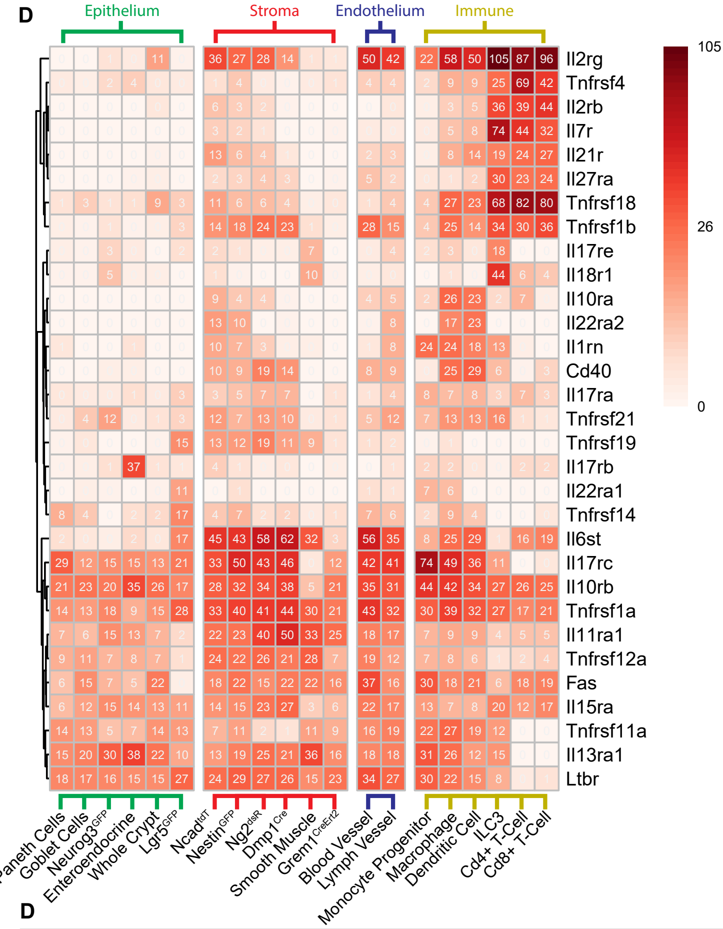


B

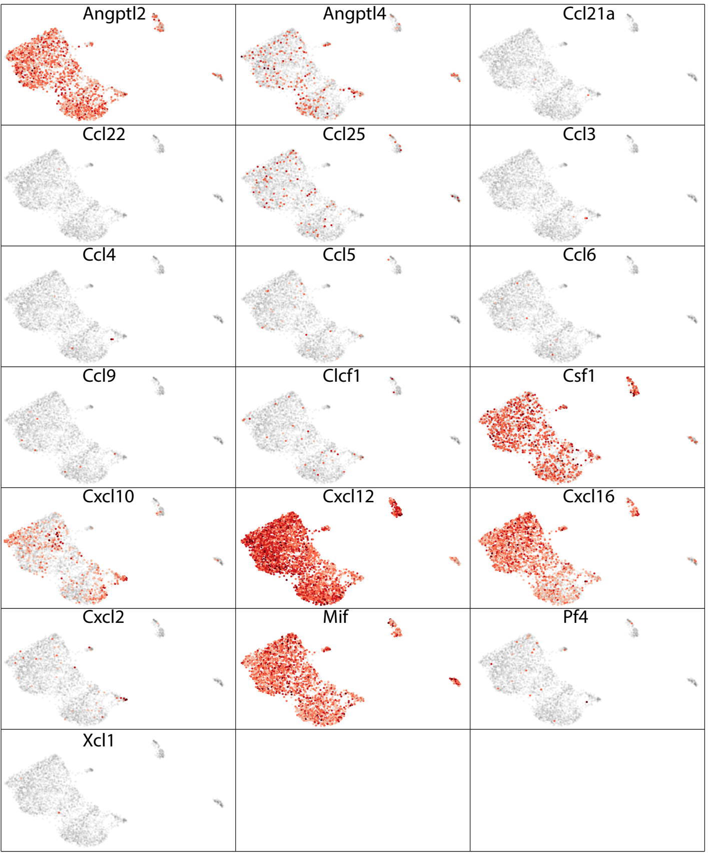


D

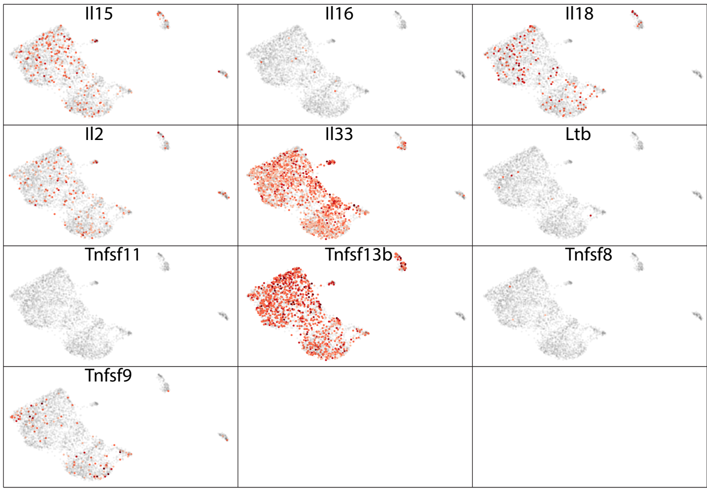

### Supplemental Figure Titles and Legends

**Figure S1. Table of Isolated Cell Types and Their Corresponding Functional genes**

(A) Table showing the list of cells used for the bRNA-seq results and the markers used in FACS isolation.
(B) Heatmap of Functional genes demonstrating the expected populations were obtained in Figure 1
